# Coastal oceanographic connectivity estimates at the global scale

**DOI:** 10.1101/2024.04.24.590881

**Authors:** Jorge Assis, Terence Legrand, Eliza Fragkopoulou, Ester A. Serrão, Miguel Araújo

## Abstract

**Motivation:** Oceanographic connectivity driven by ocean currents is critical in determining the distribution of marine biodiversity. It mediates the genetic and individual exchange between populations, from structuring dispersal barriers that promote long-term isolation to enabling long-distance dispersal that underpins species expansion and resilience against climate change. Despite its significance, comprehensive estimates of oceanographic connectivity on a global scale remain unavailable, while traditional approaches, often simplistic, fail to capture the complexity of oceanographic factors contributing to population connectivity. This gap hinders a deeper understating of species’ dispersal ecology, survival, and evolution, ultimately precluding the development of effective conservation strategies aimed at preserving marine biodiversity. To address this challenge, we present a comprehensive dataset of connectivity estimates along the world’s coastlines, known for their rich marine biodiversity. These estimates are derived from a biophysical modelling framework that combines high-resolution ocean current data with graph theory to predict multi-generational stepping-stone connectivity. Alongside, we provide coastalNet, an R package designed to streamline access, analysis, and visualization of connectivity estimates. This tool enhances the utility and application of the data, adhering to the FAIR principles of Findability, Accessibility, Interoperability, and Reusability. The dataset and package set a new benchmark for research in oceanographic connectivity, allowing a better exploration of the complex dynamics of coastal marine ecosystems.

**Main types of variables contained:** Pairwise connectivity estimates (probability and time) between coastal sites.

**Spatial location and grain:** Global, equal-area hexagons with 8.45 km edge length.

**Time period and grain:** Daily, from 2000 to 2020.

**Major taxa and level of measurement:** Coastal marine biodiversity.

**Software format:** A package of functions developed for R software.

## Introduction

Oceanographic connectivity driven by ocean currents plays a crucial role in shaping the distribution of marine biodiversity. It mediates the extent to which populations exchange genes and individuals, from structuring sharp barriers to dispersal that isolate populations and foster biodiversity differentiation, to enabling long-distance dispersal events that underpin first settlements and the expansion of species’ distributional ranges ^1–3^. Accordingly, oceanographic connectivity can facilitate the persistence of marine species in face of climate change by promoting recolonization events and improving resilience through the maintenance of genetic diversity and adaptive potential ^4,5^. The majority of marine species are dispersed by planktonic stages, with periods varying widely from a few hours to several hundred days ^6,7^. This variability further contributes to regulating population connectivity, influencing whether populations exist as isolated units or as part of a larger, interconnected metapopulation, with sources and stepping-stone processes fostering the persistence of extensive regional pools of biodiversity ^6–8^.

Oceanographic connectivity has not been well resolved at the global scale. Traditional studies often rely on simplistic methodologies, such as isolation by distance models that presuppose increased population isolation with greater geographic distances, or illustrative maps from which population connectivity is inferred from the broad-scale patterns of ocean currents ^9,10^. Such approaches significantly overlook critical oceanographic processes like gyres, tidal forces, eddies and fronts, as well as topographic features, which contribute to the creation of complex, asymmetric and highly variable connectivity flows between populations ^11,12^. Moreover, the relationship between ocean currents and population connectivity unfolds in a non-linear, skewed way ^2^, mirroring the extensive genetic structure found across marine species distributions ^13,14^. Advancements call for sophisticated biophysical models that integrate detailed oceanographic data, such as daily current velocity fields, with biological characteristics of marine species, such as spawning times and propagule periods.

By simulating the movement of particles advected by oceanographic processes, biophysical models can predict dispersal pathways between populations and final settlement locations ^15,16^. Importantly, when coupled with graph theory, biophysical models recurrently yield estimates of connectivity that mirror the observed patterns of marine biodiversity, from demography to genetics ^2,3,14,17,18^. Such an approach considers biological populations as graph vertices with their edges represented by oceanographic connectivity estimates (e.g., probability of connectivity). Specific graph algorithms can compute multigenerational stepping-stone connectivity ^12^ or identify critical connectivity components with higher centrality that can bridge populations across large water masses ^19^. Despite the significant advancements made through biophysical modeling in understanding the role of coastal oceanographic connectivity on marine biodiversity patterns ^2,3,14,20^, and its implications for marine conservation and management ^7,15,19^, research is hindered by the lack of a globally ready-to-use connectivity database. This limitation is impaired by the technical and computational requirements associated with the development of such complex modeling tools.

To address this critical data gap and unlock the full potential of research into oceanographic connectivity, we provide comprehensive connectivity estimates along the world’s coastlines, regions of critical significance owing to their rich marine biodiversity. These data are derived from a biophysical model that incorporates high-resolution oceanographic data and graph theory. Furthermore, we introduce coastalNet, an R package designed to streamline access, analysis, and visualization of coastal connectivity estimates, enhancing the utility and application of these data. Our contributions, adhering to the FAIR principles of Findability, Accessibility, Interoperability, and Reusability, set a new benchmark for research in oceanographic connectivity, allowing a better exploration of the complex dynamics of coastal marine ecosystems.

### Biophysical model

Simulations of oceanographic connectivity were conducted with a biophysical modelling framework, previously used and validated against empirical demographic and genetic data from diverse ecological groups, such as fish ^11^, molluscs ^17,18,21^, macroalgae ^3,14^, seagrass ^22^ and mangroves ^2^. The model was conducted globally with surface data on the direction and intensity of ocean currents at the daily basis, derived from the Global Ocean Physics Reanalysis (GLORYS12V1), provided by the EU Copernicus Marine Service. This is an ocean reanalysis of the global ocean at 0.08° horizontal resolution (approx. 8 km) forced by the global atmospheric reanalysis ERA-Interim and its successor the ERA5 reanalysis. The product integrates satellite and in-situ data from the Coriolis Ocean database ReAnalysis (CORA) ^23^. The accuracy of the GLORYS12V1 reanalysis in capturing oceanic processes has been validated elsewhere through comparisons with in-situ observations and other models, both at local and global scales ^24–26^.

The simulations involved the daily release of passive Lagrangian particles from the centers of hexagon-shaped coastal sites (Figure 1), each side measuring 8.45 km, over a 21-year period from 2000 to 2020. These particles were subjected to horizontal advection by the ocean velocity fields for a maximum duration of 180 days, a period that encompasses the pelagic phase of the vast majority of marine species ^7,15^. The position of each particle was calculated hourly through bilinear interpolation, in compliance with the Courant–Friedrichs–Lewy condition, until they either reached designated sink sites or got lost to the open ocean. During the simulation period, the realized connections between pair sites were recorded, alongside with the corresponding date of particle release and trajectory time. In total, the biophysical model ran for 7,670 days, releasing 204,344,140 particles from 26,642 coastal sites, resulting in 195,121,399 pairwise connectivity events.

**Figure 1.**
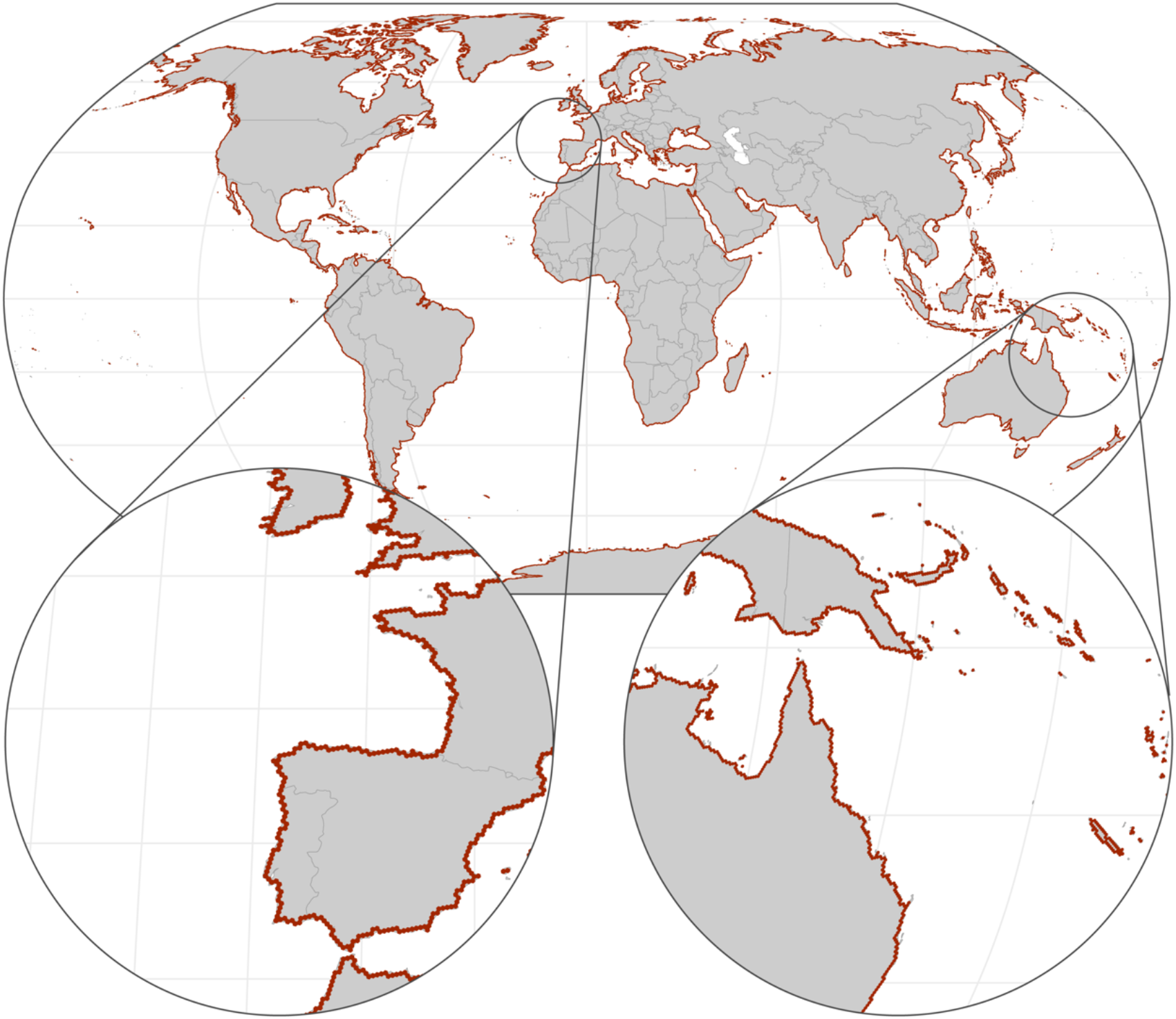
Global distribution of hexagon-shaped coastal sites (each side measuring 8.45 km) from where passive Lagrangian particles were release during the oceanographic connectivity simulations undertaken in the biophysical model.

### Oceanographic connectivity estimates

The dataset ^27^ comprises (1) a geospatial vector for geographic information systems (GIS) with the distribution of hexagon-shaped sites and the corresponding identifiers (id), and (2) a matrix of realized connections between pairs sites, with information on source site (id), sink site (id), day, month, and year of particle release and the pathway time expressed in hours.

### coastalNet R-Package

In addition to providing oceanographic connectivity estimates between pairs of sites, the coastalNet package of R functions was developed to streamline access, analysis, and visualization of connectivity estimates (refer to Table 1 for the available functions).

**Table 1.** List of functions available in coastalNet R-Package, description, user arguments, and outputs.

| Function | Description | Arguments | Output |
| --- | --- | --- | --- |
| getDataBase() | Downloads and/or loads the connectivity database. | Folder path to save the database.<br>Option to overwrite existing database. | Database containing oceanographic connectivity events. |
| getHexagonID() | Identifies hexagon IDs for a specific location or object. | Spatial object (coordinates, polygon, raster).<br>Search method (area covered, sites, centroid of sites). | List of hexagon IDs matching the search criteria. |
| getConnectivityEvents() | Extracts connectivity events from the database. | Hexagon IDs of interest.<br>Timeframe (year, month, day).<br>Duration of connectivity events (days). | Data table containing filtered connectivity events. |
| getPairwiseConnectivity() | Calculates connection strengths between pairs of sites. | Connectivity events data.<br>List of hexagon IDs to connect from.<br>List of hexagon IDs to connect to.<br>Connection metric (probability, number of events, time).<br>Option to consider stepping-stone connections. | Data tables (squared or pairs) summarizing connection strengths between sites.<br>Network representing connections between sites. |
| mapConnectivity() | Visualizes connection strengths on a map. | Connection data (from getPairwiseConnectivity). | Spatial object with lines representing connections. |

In the R environment, the getDataBase function accesses the database comprising the connectivity events between the hexagon sites. The getHexagonID function allows specifying a region of interest (e.g., study area) using various spatial data formats (e.g., spatial polygons, raster layers or matrices with georeferenced records). The getConnectivityEvents allows filtering the database, extracting the events relevant to the region of interest and timeframes, specifically, the period of particle releases (day, month, and/or year) and the particle duration period. These data are then used by the calculatePairwiseConnectivity function to generate matrices of pairwise estimates of oceanographic connectivity between sites, either as probability (forward or backward) or time estimates. For forward probability, pairwise estimates are determined by dividing the number of particle exchanged from site i to site j, by the total number of particles released by site i ^2,3,7^. For backward probability, pairwise estimates are determined by dividing the number of particle received by site i from site j, by the total number of particle received from site i ^28^. For forward time, estimates are determined by the average time of particles trajectories from site i to site j, while for backward time, estimates are determined by the average time of trajectories received in site i from site j. Finally, the mapConnectivity function allows mapping connectivity patterns.

To allow estimating asymmetrical multigeneration stepping-stone connectivity processes, the package integrates a graph-theoretical approach. To this end, the graph vertices are defined by the hexagon sites while the edges are defined by their oceanographic connectivity estimates ^7,19,29^. The Dijkstra algorithm is used to find the shortest path between sites through the reduction of the sum of log-transformed distances (e.g., -log(forward probability of connectivity ^30^). For graphs using connectivity based on probabilities, multigeneration estimates are determined by multiplying the edges along the shortest paths, while for connectivity based on time, estimates are determined by the sum along the paths. In this process, the number of stepping-stones is reported.

To install the package (during blind peer-review), the following commands should be typed into the R command prompt:

1. download.file(“https://figshare.com/ndownloader/files/45641682?private_link=d6c9f097f1bc7613029e”, destfile=“coastalNetPeerReview.zip”)
2. unzip(“coastalNetPeerReview.zip”)
3. install.packages(“devtools”)
4. devtools::install(“coastalNetPeerReview”)
5. library(coastalNet)

(This section will be removed after acceptance).

To install the package (accepted version of the manuscript), the following commands should be typed into the R command prompt:

1. install.packages(“remotes”)
2. remotes::install_github(“{hidden for blind peer-review}/coastalNet”)
3. library(coastalNet)

### Usage notes

The applications of the data provided span several pivotal fields within marine science, including ecology, evolution, conservation, and management. Among other applications, the connectivity estimates can be used to (1) explore evolutionary dynamics shaping marine biodiversity, notably by examining how ocean current patterns influence genetic differentiation across populations, (2) aid in the projection of distribution range expansions as species track suitable habitats and (3) map the potential spread of non-native species. In fisheries management, the estimates can be used to (4) evaluate the connectivity among economically significant coastal fish stocks, thereby facilitating informed decisions on harvesting rates and stock replenishment strategies, and (5) guide conservation priorities, through the identification of connectivity corridors or isolated populations in risk of disappearing. This can directly feed the design of networks of Marine Protected Areas (MPAs), where strategic planning based on connectivity can significantly support conservation efforts.

To demonstrate the application of the coastalNet package, we provide three tailored examples with supporting R code. The first example ^3^ investigates how oceanographic connectivity influences the population differentiation of a kelp species (*Laminaria ochroleuca*). The code builds a statistical regression model fitting the connectivity estimates against empirical genetic differentiation data and maps stepping-stone connectivity between populations (Figure 2a; Supplement 1). Overall, the example shows that oceanographic connectivity influences population differentiation of the kelp species, explaining ∼65 % of genetic data variability. The second example ^19^ maps oceanographic connectivity of fish populations along a network of Mediterranean MPAs. The code estimates fish connectivity (using a typical pelagic larvae duration of 32 days^7,15^) between spatial polygon data representing the shape and distribution of MPAs (Figure 2b; Supplement 2). Overall, the example shows vast network connectivity along the western Mediterranean Sea. This contrasts with the eastern Mediterranean Sea, where insufficient stepping-stone connectivity was inferred between MPAs, likely precluding proper conservation of regional pools of fish biodiversity in the long term. The third example explores how oceanographic connectivity may disrupt the ability of a kelp species (*Macrocystis pyrifera*) to reach suitable habitats while shifting ranges under future climate change. The code combines the connectivity estimates with habitat suitability data for the present and the future ^31^. Overall, it identifies vast areas with suitable habitats in the future that might not be colonized due to dispersal barriers structured by ocean currents (Figure 2c; Supplement 3).

**Figure 2.**
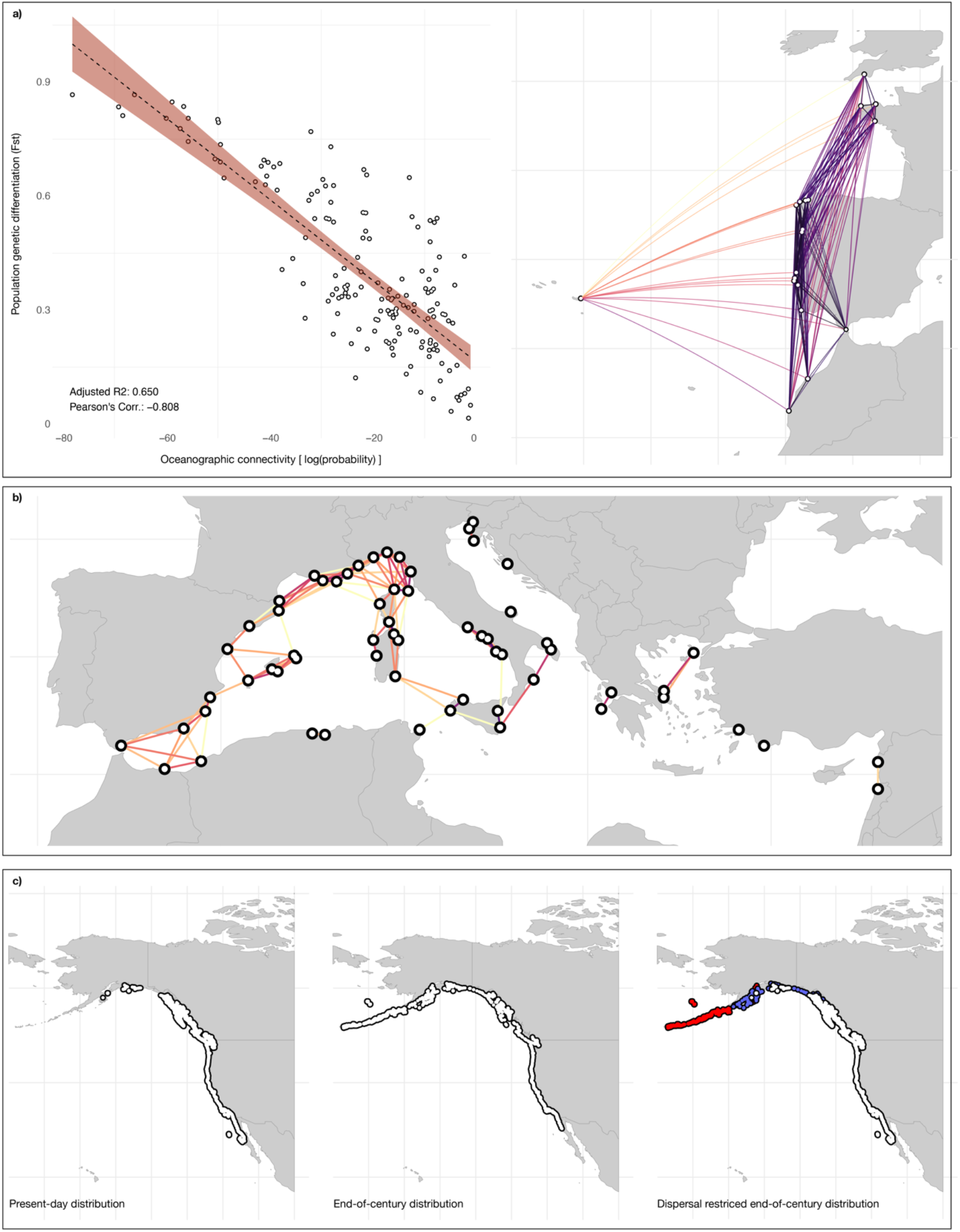
Demonstrative outputs of coastalNet R package (tailored examples with supporting R code in Supplements 1-3). (panel a) Exploring how oceanographic connectivity influences the population differentiation of a kelp species (*Laminaria ochroleuca*). (panel b) Mapping oceanographic connectivity of fish populations along a network of Mediterranean MPAs. (panel c) Exploring how oceanographic connectivity may disrupt the ability of a kelp species (*Macrocystis pyrifera*) to reach suitable habitats while shifting ranges under future climate change.

## Supplementary information

Supplementary information 1. Demonstrative output of coastalNet R package with R code exploring how oceanographic connectivity influences the population differentiation of a kelp species (*Laminaria ochroleuca*).

Supplementary information 2. Demonstrative output of coastalNet R package with R code mapping oceanographic connectivity of fish populations along a network of Mediterranean MPAs.

Supplementary information 3. Demonstrative output of coastalNet R package with R code exploring how oceanographic connectivity may disrupt the ability of a kelp species (*Macrocystis pyrifera*) to reach suitable habitats while shifting ranges under future climate change.

## Data availability statement

The geospatial vector for geographic information systems with the distribution of hexagon-shaped sites and the matrix of realized connections between pairs sites are temporarily available at: https://figshare.com/s/d6c9f097f1bc7613029e (the URL of the final repository will be provided after acceptance).

## Code availability

The source code and documentation of the coastalNet package are temporarily available at: https://figshare.com/ndownloader/files/45641682?private_link=d6c9f097f1bc7613029e (the URL of the final repository will be provided after acceptance).

## Competing interests

The authors declared no conflict of interest.

## Notes

https://figshare.com/s/d6c9f097f1bc7613029e

### Competing Interest Statement

The authors have declared no competing interest.

